# transcranial Direct Current Stimulation Studies Open Database (tDCS-OD)

**DOI:** 10.1101/369215

**Authors:** Pnina Grossman, Ivan Alekseichuk, Gabriel de Lara, Kunal Paneri, Patrik Kunz, Zsolt Turi, Marom Bikson, Walter Paulus, Andrea Antal

## Abstract

Transcranial direct current stimulation (tDCS) is increasingly evaluated for a myriad of clinical indications and performance enhancement mainly related to cognitive and motor applications. As a result tDCS has been tested with diverse inclusion criteria and subject populations. tDCS is customized for applications by many factors, including stimulation dose (montage, current, time), associated training and subject inclusion/exclusion criteria. The rapid proliferation of applications, technological advancements and emerging scientific discoveries presents challenges to the organization and consolidation of tDCS data – which can lead to scientific, clinical and public confusions. In this paper, we develop a system to summarize and consolidate methodological aspects of tDCS, concentrating on study design and the physical parameters of the stimulation. We introduce a community-driven, open access database, where these parameters, including stimulation intensities, duration, electrode sizes, montages, and subject information, are noted for relevant tDCS publications. The transcranial Direct Current Stimulation Studies Open Database (tDCS-OD) (http://tdcsdatabase.com) will support constructive dialogue on research and clinical tDCS applications, critical evaluation of past work, and identification of promising protocols, while reducing ambiguity about stimulation methodology. The database design allows ongoing updates and editing, with transparent version control. At the moment, the database includes information about 56,613 tDCS sessions.

## Introduction

Non-invasive neuromodulation using transcranial direct current stimulation (tDCS) provides researchers and medical scientists with a tool to probe and modify brain functions to treat neuropsychiatric conditions. Due to its apparent simplicity [1], tolerability [2];[3], accessibility [4], safety [2] and potential benefits, the use of tDCS is further increasing. While until 2005, the aggregated number of published tDCS papers was ~300, a decade later ~3000 articles have been published from teams around the globe. The increase in study volume and breadth went along with a large and diverse volume of published tDCS trials [2];[5];[6];[7];[8];[9]. The proliferation of tDCS studies has created challenges for aggregating data, but also an opportunity to consolidate experience.

Over a decade of research efforts have supported mechanistic understanding of tDCS, optimized stimulation protocols, and improved reproducibility and standardization [10];[2];[11]. Nevertheless, alongside accelerated interest in tDCS, many scientific and methodological caveats have emerged, which engender closer examination of long-standing assumptions and suggest the need for a more nuanced trial design [12];[13];[14]. tDCS has a parameter space of dose, with the behavioral task and inclusion criterion increasing dimensionality. Precisely because tDCS outcomes are considered nuanced to protocol design, the considerable variation in stimulation parameters and protocols poses a challenge to the identification of the most promising protocols and convergence upon standard methodologies.

To help address this problem, we introduce here a new, open access community-driven database (http://tdcsdatabase.com). There the stimulation parameters, including the intensity, duration, electrode size, montage, number of participants/patients etc., are tabulated – supporting study identification and meta-analysis across research studies meeting specific trial design criterion. Precise cataloging of past methods underpins review, replication, and design of tDCS trials. We encourage scientists to use the database and to further improve its efficacy by editing and correcting tentative errors. Online instructions are available. As a transparent and collective effort, this database will support ongoing rigorous advancement of the field.

## Methods

The methods herein described serve to 1) explain the structure of the database (i.e. parameters indexed for each publication); 2) our process of data aggregation as a detailed guidance document for ongoing contributions from members of the scientific community in updating the online database; and 3) outline our methods for initially populating the database with some illustrative analysis.

### Database structure and population

A python script was created to search the PubMed database for the keywords “transcranial direct current stimulation”. Basic information about the paper (e.g. title, authors, publication date) was automatically extracted by script and entered into the database. Strict inclusion criteria for the tDCS database are published scientific papers (or any format including original articles, review, editorials, case-reports, book chapters) with a unique PMID or ISBN identifier.

The database will be repopulated monthly, on the first day of each month. To do so, the python script will search PubMed for any articles published between the previous update and the date on which the database is being repopulated. As the tDCS Open Database may be used in subsequently (published) analysis, the database will include monthly “save points.” On the first of each month, before the database is updated, the website will create a permanent record of what was in the database at that time. These records can then be checked against any publications that cite the database to verify the complete data included at the time of citation.

### Data aggregation process

An automatic search of databases for the keywords chosen returns many articles not relevant for the tDCS database. A “relevance” category is included for each paper with three options: “relevant”, “not relevant”, or “relevance unspecified”. New papers that are automatically added will be categorized as “relevance unspecified” by default. Papers that, upon examination, are found to be not relevant for the purpose of the database (i.e. do not meet the inclusion criteria given below) are not deleted, as they could then be re-added later by python script update function or manually by a user. And indeed, the record and rational for the study exclusion from subsequent analysis should be saved. Instead, these irrelevant articles are kept in the system but marked as “not relevant”. Marking a paper as “not relevant” requires the editor to input a reason that a paper is not relevant e.g. “Review without new session data”, “Not tDCS”, “Not peer reviewed scientific paper”, “Not human subjects”.

The criteria for papers to be marked “relevant” were as follows: papers needed to have (1) used tDCS, (2) been peer reviewed, (3) been tested on human subjects, (4) reported at least one original session, (5) reported all or meaningful details on dosage.

Papers that meet these criteria for relevance are manually analyzed by editors of the website to extract relevant data from them. These data include information on each experiment performed using tDCS. The information on experimental procedure includes the current at which tDCS was administered (measured in milliamperes), the duration of the tDCS session (measured in minutes), the electrode sizes (measured in cm^2^), and positions of each electrode (in EEG 10-20 format where applicable) with an indication of what type (nominally active, nominally return, unspecified or other) the authors considered each electrode. Variations in any of these four fields are considered and noted as separate experiments within the same study, even if both experiments are in the same subject group.

For each experiment, demographic information about the subjects is included, provided that this information was in the paper. The demographic information includes the number of subjects in each experiment, their gender, and their age, measured by average and standard deviation, and/or range. To account for the fact that some papers use the same subjects in different experiments and some use entirely different subjects, the total number of subjects and their gender is also collected, as well as the total number of subjects with any medical condition. In the case that demographic information was only provided for all subjects, it is assumed that this information can be applied to subjects in individual experiments. It is also noted for each experiment whether there was a control group who received sham tDCS, as well as if there were any subjects in repeated sessions, defined as more than 4 sessions in a given week, and, if so, how many weeks of repeated sessions there were. This threshold for repeated is arbitrary, and future versions of the database will make this threshold selectable in search. A total number of sessions is calculated for each experiment by taking every subject who received non-sham tDCS and multiplying this by the number of non-sham tDCS sessions they received.

Relevant papers must include at least some new tDCS session data. For example, a meta-analysis of prior studies which includes no previously unpublished data would not be marked “relevant”. However, a review which has some (e.g. 1 session or more) original data can be marked “relevant”, with commentary as needed on the selection of only new sessions. Any commentary should clearly explain the logic used to determine which sessions are new vs overlapping with prior publications in the effort to ensure only session not already counted in the database in prior publications are counted for the review. A paper that aggregates tolerability results across studies would only be marked “relevant” for new subjects not otherwise reported, meaning a tolerability review that only summarizes data collected from subjects including in prior publications would not be scored as including any new subjects. Still, in such cases, it is again useful to use the comment section to explain this decision and moreover to list the original. Thus, comments on papers considered “not relevant” provide important content to the database record.

### Website structure

The database can be accessed at tdcsdatabase.com. Accessing the database and downloading all content does not require registration or an account. For editing, users will register with their email address (which is not public) and choose a publicly visible user name. Once registered and logged in, the user can either submit edits for an existing entry, or add a new entry. All changes must be approved by a website administrator before they are accepted into the database. Once accepted, the entry will show that it was edited by a specific user on a specific date. The users can also access an account section of the website where they can edit their user information, and see if their edits to any entry were accepted by an administrator, and, if not, why the edits were not accepted. The website will feature step-by-step tutorials for how to correctly collect data from an article an entry on the website, as well as information callouts. A summary of how information should be collected is presented in Table 1 (with significantly more detailed and updated instructions available online).

**Table 1:** Abridged Instructions for Entering Article Data into Database

| Field | Instructions for filling out field |
| --- | --- |
| PMID | PMID given by the PubMed database. Once entered, this can be used to Auto-Populate the Title, Abstract, Authors, Year Published, DOI and Journal Keywords fields. If an article does not have a PMID, the No PMID button must be pressed to fill out any of those fields manually. |
| Title | Title of article. Can be filled out manually or using Auto Populate with a valid PMID. |
| Abstract | Abstract of article. Can be filled out manually or using Auto Populate with a valid PMID. |
| Authors | Authors of article, listed in the format "Last name, First initial". Can be filled out manually or using Auto Populate with a valid PMID. |
| Year Published | Year article was published. Can be filled out manually or using Auto Populate with a valid PMID. |
| DOI | DOI. Can be filled out manually or using Auto Populate with a valid PMID. If article does not have a PMID, this field will be mandatory. |
| Journal Keywords | Keywords given on PubMed. Can be filled out manually or using Auto Populate with a valid PMID. |
| Keywords | Any additional keywords the user wants to apply to the article. |
| Relevant for database | Determination of whether an article is relevant for the database. New articles will default to "unspecified." If examination of article reveals that article is relevant (papers needed to have (1) used tDCS, (2) been peer reviewed, (3) been tested on human subjects, (4) reported original sessions, (5) reported "meaningful" details on dosage) select relevant. If examination of article reveals that article is not relevant, select "not relevant" and add reason in "Private Note to Admin." |
| Reason Article Not Relevant | Reason for not including article in database if article is deemed not relevant by editor. |
| Total # of Subjects in Paper | Total number of subjects who received tDCS sessions in paper. Do not count subjects twice if they participated in two separate experiments. Do not count subjects if they only received sham tDCS or were part of a control group that did not receive tDCS. |
| Total # of Females in Paper | Total number of female subjects who received tDCS sessions in paper. Do not count female subjects twice if they participated in two separate experiments. Do not count female subjects if they only received sham tDCS or were part of a control group that did not receive tDCS. |
| Subjects with Medical Condition | Total of number of subjects with a given medical condition. If a medical condition is not listed, fill out condition under other. Do not count subjects twice if they participated in two separate experiments. Do not count subjects if they only received sham tDCS or were part of a control group that did not receive tDCS. |
| Experiment #1 | Conditions under which subjects received tDCS sessions. If current, duration, electrode placement or electrode size varied, add another experiment. All relevant papers must have at least one Experiment. |
| Current (mA) | The current at which non-sham tDCS sessions were conducted for this experiment, given in milliamperes. There is no need to indicate if current was ramped-up/ramped-down. |
| Duration (min) | The duration of non-sham tDCS sessions for this experiment, given in minutes. Do not count the ramp-up/ramp-down time. |
| Total Sessions | The number of sessions in an experiment is given by multiplying the number of subjects who received non-sham tDCS stimulations by the number of non-sham tDCS stimulations they received. |
| # of Subjects in Experiment | Number of subjects who received non-sham tDCS sessions in this experiment. If a subject participated in more experiment, count subject in this field for each experiment they participated in. |
| # of Females in Experiment | Number of female subjects who received non-sham tDCS sessions in this experiment. If a female subject participated in more experiment, count subject in this field for each experiment they participated in. If only a total number of female subjects for the paper was given, assume an even distribution over all experiments, using only whole numbers of female subjects. |
| Age (avg) | Average age of subjects in an experiment, given in years. If average age of all subjects in paper is given, but not the average age of subjects in each experiment, assume an even distribution of ages. |
| Age (SD) | Standard deviation of ages of subjects in an experiment, given in years. If standard deviation of age of all subjects in paper is given, but not the standard deviation of age of subjects in each experiment, assume an even distribution of ages. |
| Age range (min) | Minimum age of subjects in experiment, given in years. If age range minimum was given for all subjects, but not each experiment, use age range minimum for all subjects for each experiment |

| **Field** | **Instructions for filling out field** |
| --- | --- |
| Age range (max) | Maximum age of subjects in experiment, given in years. If age range maximum was given for all subjects, but not each experiment, use age range maximum for all subjects for each experiment |
| Electrode #1 | Each Experiment must have a minimum of 2 electrodes. Additional electrodes can be added using the "Add Electrode" button. Position, Size and Authors Consider fields will appear for each added electrode. |
| Electrode #1 Position | Position of Electrode #1, using EEG 10-20 system, if possible. |
| Electrode #1 Size | Size of Electrode #1, given in cm<sup>2</sup>. |
| Authors Consider Electrode #1 | The type of electrode, as described by the authors of the article, is selected from the options "Nominally Active", "Nominally Return", "Other" or "Unspecified". |
| Electrode #2 | Each Experiment must have a minimum of 2 electrodes. Additional electrodes can be added using the "Add Electrode" button. Position, Size and Authors Consider fields will appear for each added electrode. |
| Electrode #2 Position | Position of Electrode #2, using EEG 10-20 system, if possible. |
| Electrode #2 Size | Size of Electrode #2, given in cm<sup>2</sup>. |
| Authors Consider Electrode #2 | The type of electrode, as described by the authors of the article, is selected from the options "Nominally Active", "Nominally Return", "Other" or "Unspecified". |
| Sham controlled? | Whether an experiment had a sham control group. |
| # Subjects in repeated sessions | Number of subjects in an experiment who received more than 4 non-sham tDCS sessions in a week. |
| # Weeks of repeated sessions | Number of weeks that each subject received repeated tDCS sessions. |
| Notes | Any additional notes that could be important for the website administrators or other researchers. |

Without need to register, researchers can use the website to look up information in the database. Individual articles can be searched for by PMID, title, or author. Groups of articles can also be filtered by field (e.g. current, duration, etc.) as well as demographic information and year published. The articles returned for these filters by default will be those marked as relevant and relevance unspecified, but this too can be adjusted by the researcher. The researcher can download the data from the website in standard format, such as an Excel. Work leveraging the website should cite the website (with date of search) and this article.

### Initial database manual population

Papers published before July 1, 2013 were manually data-mined by one investigator, went through a quality control check by at least one other investigator, with any inconsistencies resolved by consensus of authors – with a total of 32,580 sessions extracted from this period [3]. Papers published on or after July 1, 2013 and before Jan 1, 2016 were manually data-mined by a researcher, but the results were not independently verified except a superficial review for consistency and outliers– with a total of 24,033 sessions scored from this period. Papers on or after Jan 1, 2016 and before Jan 1, 2018 were indexed by our automatic search but not data mined, thus papers from this period were marked as “relevance unspecified”. Given that the goal of this effort was to design an open database which support ongoing editing by the community, we consider this existing substantial manual data-mining by the co-authors sufficient to: 1) establish a critical mass to encourage ongoing data base use and editing by the scientific community; and 2) identify the suitability of our relevance marking system and database design.

In addition to the general data base inclusion for papers, for only the purposes of our initial manual analysis we applied additional inclusion criterion: (1) used an electrolyte-rich contact material, (2) clearly (unambiguously) reported dosage information and (3) were published in English. These are in addition to inclusion criteria for the database in general.

## Results

At the time of this paper, the automatic data-based population (conducted on Jan 1, 2018, and including papers published before Jan 1, 2018) captured 4,713 total papers, the amount of papers published per year can be seen in Figure 1. General information on these papers (PMID; Title; Abstract; Authors; Year Published; DOI; Journal Keywords) was extracted and populated into the tDCS Open Database (tdcsdatabase.com). These papers do not necessarily meet the relevance criteria for the database; relevance will be determined when the papers are manually reviewed on the website.

**Figure 1:**
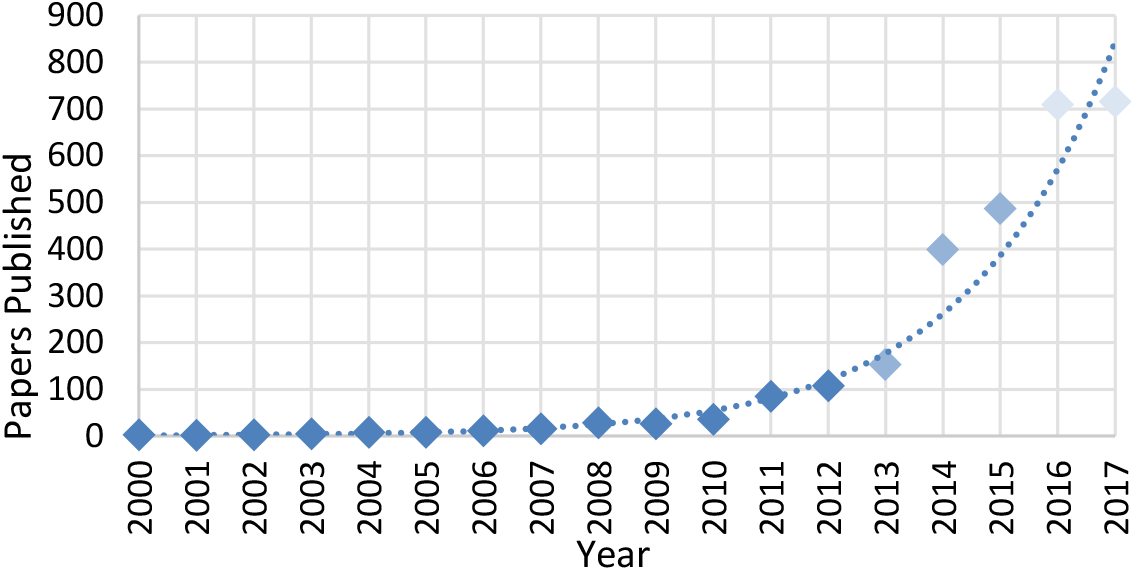
Papers published per annum that included tDCS sessions which matched the criteria listed as of January 3^rd^, 2018. While this establishes an exponential growth trend, it should be noted that the number of papers published after July 1^st^, 2013 (as indicated with a lighter shade of blue) were not independently verified to have met the inclusion criteria, and the number of papers published in 2016 and 2017 (as indicated with the lightest shade of blue) were not sorted for relevance. An exponential trend line based on the data up to 2015 was added.

Our initial manual analysis for papers published before Jan 1, 2016 and meeting further inclusion criterion (see Methods). From a total of 1,325 papers automatically extracted and given an initial relevance grading for this period, session information from 809 papers was manually entered into the tDCS Open Database. This initial manual aggregation included 56,613 sessions applied across >20,000 subjects. Basic application, dose, and demographic information as reflected in the current data set are shown (Figures 2-4).

**Figure 2:**
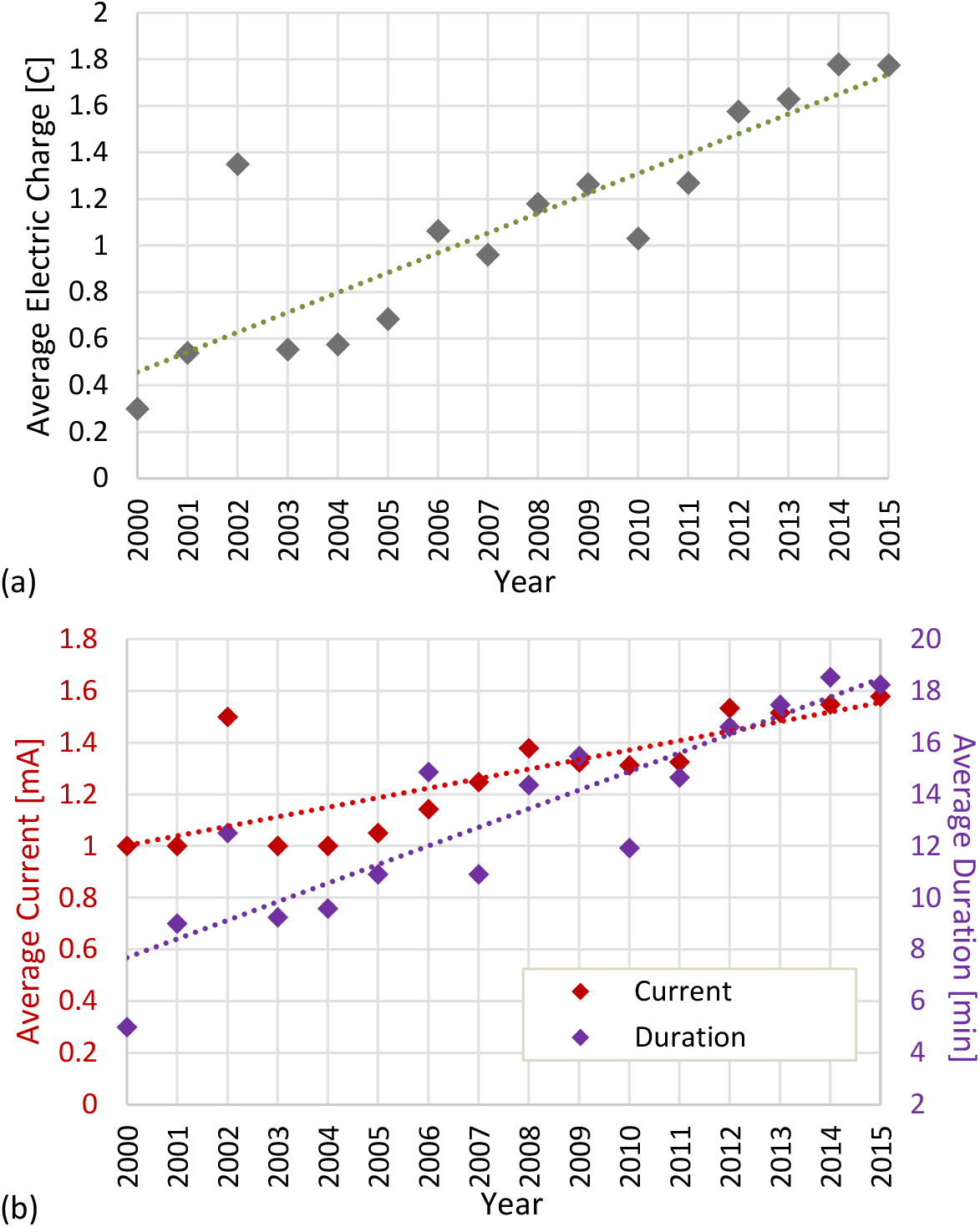
Average Electric Charge (a) and Average Current/Duration (b) vs. Year. Each experimental set-up that gave information on the current and duration was evenly weighted in calculating averages. A linear trend line was added. All data are from articles published in or before 2015 and accessed on February 5^th^, 2018.

**Figure 3:**
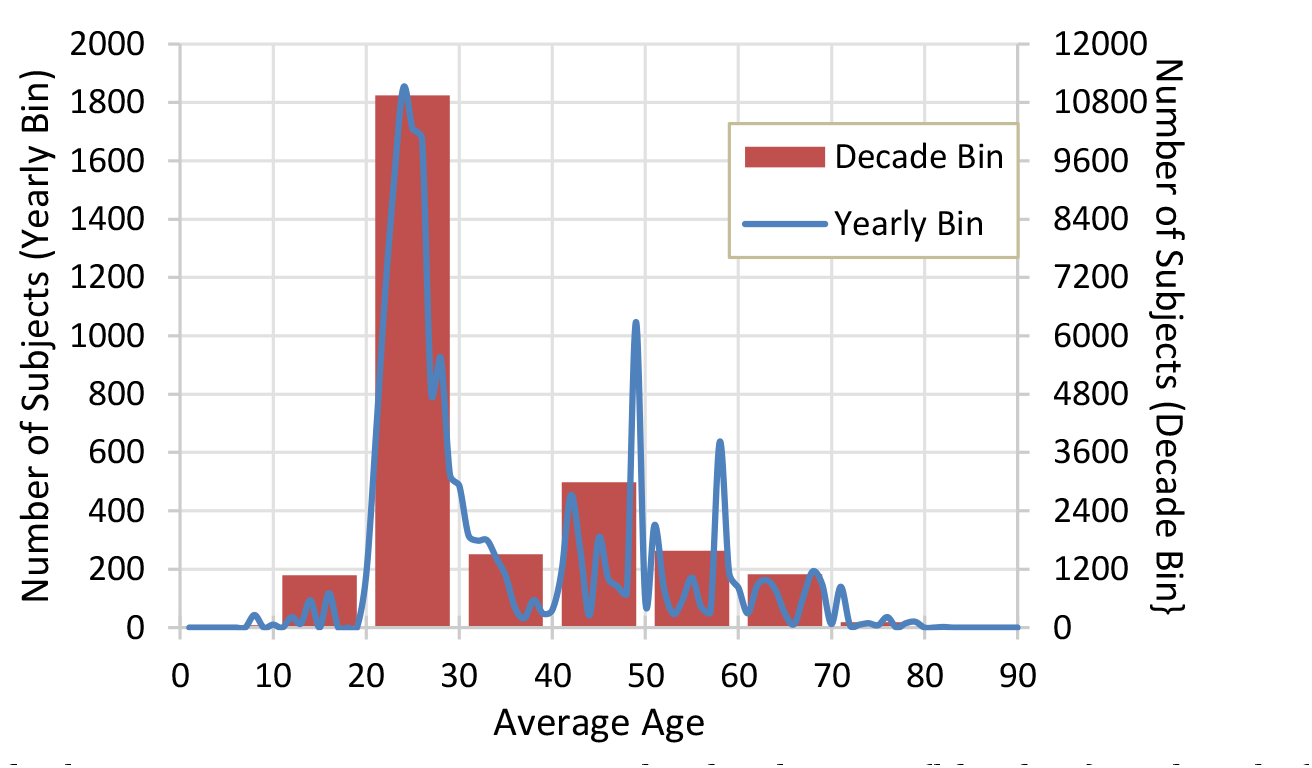
Number of subjects at an average age, grouped either by year (blue line) or decade (orange bar). If average age was provided, it was applied to all subjects in an experiment. If no average age was provided, the mid-point of age range was used instead. All data are from articles published in or before 2015 and accessed on February 5^th^, 2018.

**Figure 4:**
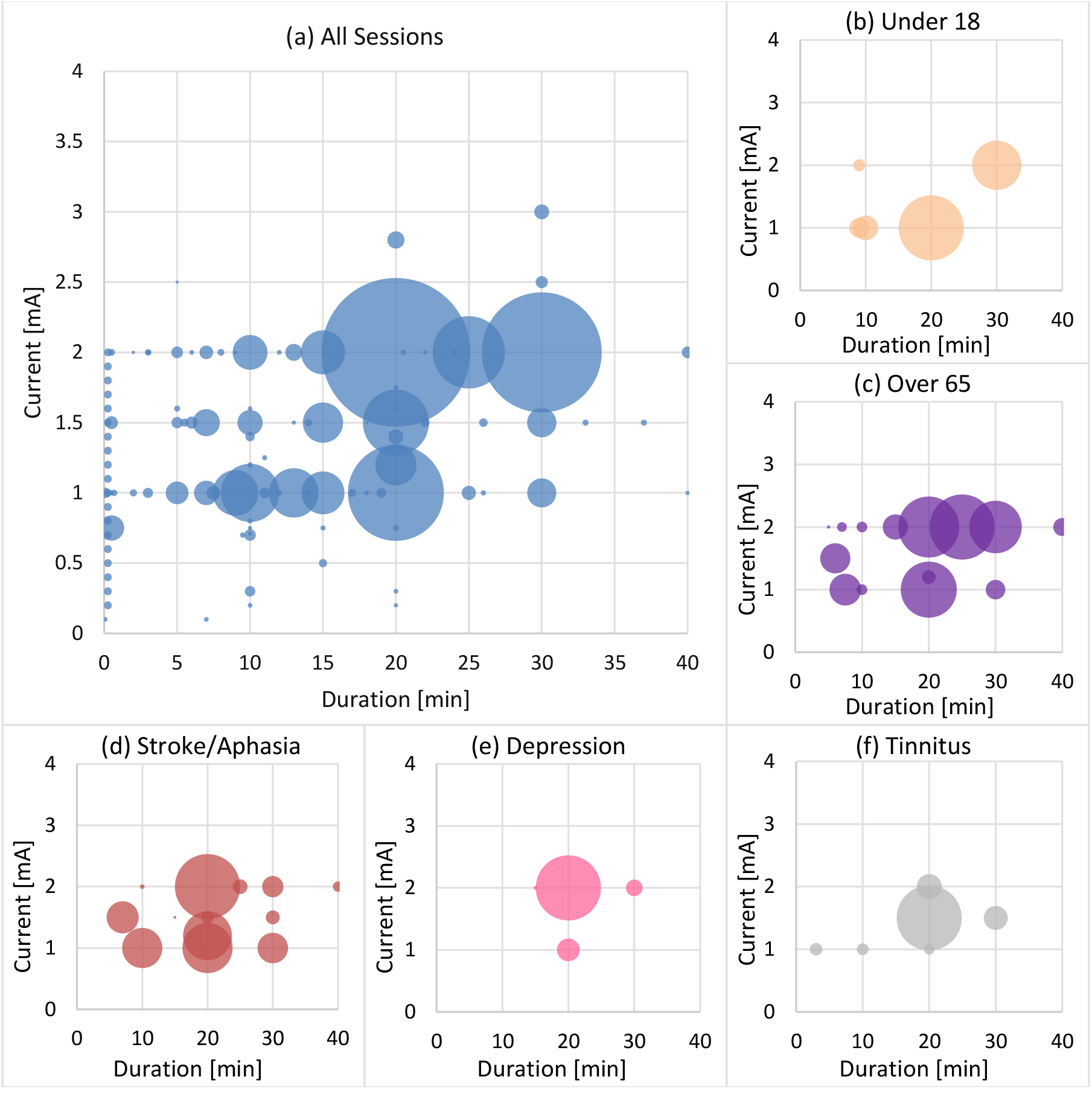
Heat map of tDCS sessions grouped by current and duration for all sessions (a), under 18 (b), over 65 (c), stroke/aphasia (d), depression (e), tinnitus (f). Ages were established using the average age protocol from Figure 3. Stroke/aphasia, depression and tinnitus were chosen because they were the three medical conditions with the highest number of subjects that had undergone tDCS. All data are from articles published in or before 2015 and accessed on February 5^th^, 2018.

## Discussion

The tDCS Open Database goes beyond mere listing of literature search as provided e.g. by PubMed. It provides a structure for searching and combined searching of the key parameters used in tDCS for free without the necessity for individual registration. The portal allows also editing of content by any registered user with record of changes and version control. The advantage is certainly that many contributors can contribute more details as a small editing group, the disadvantage of course no editing of new contributions such as in a “Wikipedia-like” structure. Edits require approval by an administrator to preserve website integrity (preventing sloppy or malicious additions) and ensure that the data is consistent with the tDCS Open Database guidance (preventing errant fields). We hope database users will revisit entries on an ongoing basic to provide continued quality control. We encourage authors of tDCS studies to search for their articles and contribute to the database by manually editing and controlling the quality of their own article in the database. While allowing for regular updates, version control with complete historical records allows the tDCS Open Database to serve rigorous scholarship (e.g. meta-analysis). Document citing the tDCS Open Database should do so as noting the data of search and data extraction.

The categories and method of indexing tDCS papers were developed by the authors with the goal to balance the completeness and itemization of the summary protocols from thousands of papers. Inevitably, situations will arise where a paper, or combination of papers, presents a conundrum as far as how to complete the existing summary fields. The open comment section for each paper allows at least for a noting of such concerns.

The tDCS Open Database (tDCS-OD) includes embedded tools to filter and extract data, as well as a simple export function to analyze and display data using independent software. Later versions may include embedded data processing and plotting functions. As this code for the tDCS Open Data is itself open source, we invite collective efforts to enhance these tools.

## Acknowledgement

We thank Vincent Lubbe for his help during the data review process.

